# Robust Modelling of Additive and Non-additive Variation with Intuitive Inclusion of Expert Knowledge

**DOI:** 10.1101/2020.04.01.019497

**Authors:** Ingeborg Gullikstad Hem, Maria Lie Selle, Gregor Gorjanc, Geir-Arne Fuglstad, Andrea Riebler

## Abstract

We propose a novel Bayesian approach that robustifies genomic modelling by leveraging expert knowledge through prior distributions. The central component is the hierarchical decomposition of phenotypic variation into additive and non-additive genetic variation, which leads to an intuitive model parameterization that can be visualised as a tree. The edges of the tree represent ratios of variances, for example broad-sense heritability, which are quantities for which expert knowledge is natural to exist. Penalized complexity priors are defined for all edges of the tree in a bottom-up procedure that respects the model structure and incorporates expert knowledge through all levels. We investigate models with different sources of variation and compare the performance of different priors implementing varying amounts of expert knowledge in the context of plant breeding. A simulation study shows that the proposed priors implementing expert knowledge improve the robustness of genomic modelling and the selection of the genetically best individuals in a breeding program. We observe this improvement in both variety selection on genetic values and parent selection on additive values; the variety selection benefited the most. In a real case study expert knowledge increases phenotype prediction accuracy for cases in which the standard maximum likelihood approach did not find optimal estimates for the variance components. Finally, we discuss the importance of expert knowledge priors for genomic modelling and breeding, and point to future research areas of easy-to-use and parsimonious priors in genomic modelling.

## Introduction

Plant breeding programs are improving productivity of a range of crops and with this addressing the global and rising hunger problem that impacts 820 million people across the world (FAO *et al.*, 2019). One of the most important food sources in the world is wheat (Shewry and Hey, 2015), however, recent improvements in wheat yield are smaller than the projected requirements (Ray *et al.*, 2013) and might become more variable or even decrease due to climate change (Asseng *et al.*, 2015). This trend is in stark contrast to the United Nation’s Sustainable Development Goals that aim to end hunger and malnutrition by 2030 (General Assemby of the United Nations, 2015). Breeding has contributed significantly to the improvement of wheat yields in the past (e.g., Mackay *et al.*, 2011; Rife *et al.*, 2019), and the recent adoption of genomic selection could enable further significant improvements (Gaynor *et al.*, 2017; Belamkar *et al.*, 2018; Sweeney *et al.*, 2019).

Breeding programs generate and evaluate new genotypes with the aim to improve key characteristics such as plant height, disease resistance and yield. Nowadays, a key component in breeding is genomic modelling, where we aim to reduce environmental noise in phenotypic observations and associate the remaining variation with variation in individual genomes. We use these associations to estimate genetic values for phenotyped or even non-phenotyped individuals and with this identify the genetically best individuals (Meuwissen *et al.*, 2001). Improving this process involves improving the methods for disentangling genetic variation from environmental variation.

Genetic variation can be decomposed into additive and non-additive components (Fisher, 1918; Falconer and Mackay, 1996; Lynch *et al.*, 1998; Mäki-Tanila and Hill, 2014). Additive variation is defined as variation of additive values, which are sums of allele substitution effects over the unobserved genotypes of causal loci. Statistically, the allele substitution effects are coefficients of multiple linear regression of phenotypic values on causal genotypes coded in an additive manner. Non-additive variation is defined as the remaining genetic variation not captured by the additive values. It is commonly partitioned into dominance and epistasis values. Dominance values capture deviations from additive values at individual loci. Epistasis values capture deviations from additive and dominance values at combinations of loci. Statistically, dominance and epistasis values capture deviations due to allele interactions at individual loci and combinations of loci, respectively. Modelling interactions between two loci at a time gives additive-by-additive, additive-by-dominance and dominance-by-dominance epistasis. Modelling interactions between a larger number of loci increases the number of interactions.

Estimates of genetic values and their additive and non-additive components have different applications in breeding (Acquaah, 2009). Breeders use estimates of additive values to identify parents of the next generation, because additive values indicate the expected change in mean genetic value in the next generation under the assumption that allele frequencies will not change. Breeders use estimates of genetic values to identify individuals for commercial production, because genetic values indicate the expected phenotypic value. Estimates of genetic values are particularly valuable in plant breeding where individual genotypes can be effectively cloned. While genomic modelling currently focuses on additive values (Meuwissen *et al.*, 2001; Varona *et al.*, 2018), the literature on modelling non-additive variation is growing (Oakey *et al.*, 2006; Wittenburg *et al.*, 2011; Muñoz *et al.*, 2014; Bouvet *et al.*, 2016; Martini *et al.*, 2017; Vitezica *et al.*, 2017; Varona *et al.*, 2018; de Almeida Filho *et al.*, 2019; Santantonio *et al.*, 2019; Tolhurst *et al.*, 2019; Martini *et al.*, 2020). Notably, modelling non-additive variation has been shown to improve the estimation of additive values in certain cases (Varona *et al.*, 2018).

However, modelling non-additive variation is challenging because it is difficult to separate non-additive variation from additive and environmental variation even when large datasets are available (e.g., Misztal, 1997; Zhu *et al.*, 2015; de los Campos *et al.*, 2019). Further, pervasive linkage and linkage disequilibrium are challenging the decomposition of genetic variance into its components (Gianola *et al.*, 2013; Morota *et al.*, 2014; Morota and Gianola, 2014). This suggests that genomic modelling needs *robust* methods that do not estimate spurious non-additive values and whose inference is *stable* for all sample sizes.

One way to handle partially confounded sources of variation is to take advantage of expert knowledge on their absolute or relative sizes. Information about the relative magnitude of the sources of phenotypic variation has been collated since the seminal work of Fisher (1918). The magnitude of genetic variation for a range of traits is well known (e.g., Houle, 1992; Falconer and Mackay, 1996; Lynch *et al.*, 1998). Data and theory indicate that the majority of genetic variation is captured by additive values (Hill *et al.*, 2008; Mäki-Tanila and Hill, 2014), while the magnitude of variation in dominance and epistasis values varies considerably due to a range of factors. For example, there is no dominance variation between inbred individuals by definition. Further, model specification has a strong effect on the resulting estimates (e.g., Huang and Mackay, 2016; Martini *et al.*, 2017; Vitezica *et al.*, 2017; Martini *et al.*, 2020). With common model specifications, additive values capture most of the genetic variation because they capture the main effects (in the statistical sense), while dominance and epistasis values capture interaction deviations from the main effects (Hill *et al.*, 2008; Mäki-Tanila and Hill, 2014; Hill and Mäki-Tanila, 2015; Huang and Mackay, 2016).

In a Bayesian setting we can take advantage of such expert knowledge through prior distributions; see Gianola and Fernando (1986); Sorensen and Gianola (2007) for an introduction to Bayesian methods in animal breeding and quantitative genetics, respectively. Prior distributions reflect beliefs and uncertainties about unknown quantities of a model and should be elicited from an expert in the field of interest (O’Hagan *et al.*, 2006; Farrow, 2013). Intuitively, prior distributions allow expert knowledge to act as additional observations, and make the current analysis more robust by borrowing strength from past analyses. Priors can improve weak identifiability of the sources of variation by guiding inference towards expert knowledge when the information in the sample is low. However, quantification of the effective number of samples added by a prior is only available in specific situations (Morita *et al.*, 2008).

We propose an easy-to-use, intuitive, and robust Bayesian approach that builds on two recent innovations in Bayesian statistics: 1) the hierarchical decomposition prior framework (Fuglstad *et al.*, 2020) to provide an hierarchical description of the decomposition of phenotypic variation into different types of variation, and 2) the penalized complexity prior framework (Simpson *et al.*, 2017) to facilitate robust genomic modelling. The key ideas of the approach are that (i) visualization eases model specification and communication about the model (see Figure 1), (ii) hierarchical decomposition of variation makes it easy to incorporate expert knowledge on e.g. heritability, (iii) leveraging expert knowledge provides robust methods, and (iv) comparison of posterior distributions and prior distributions reveal the amount of information the data provided on the decomposition of variation.

**Figure 1:**
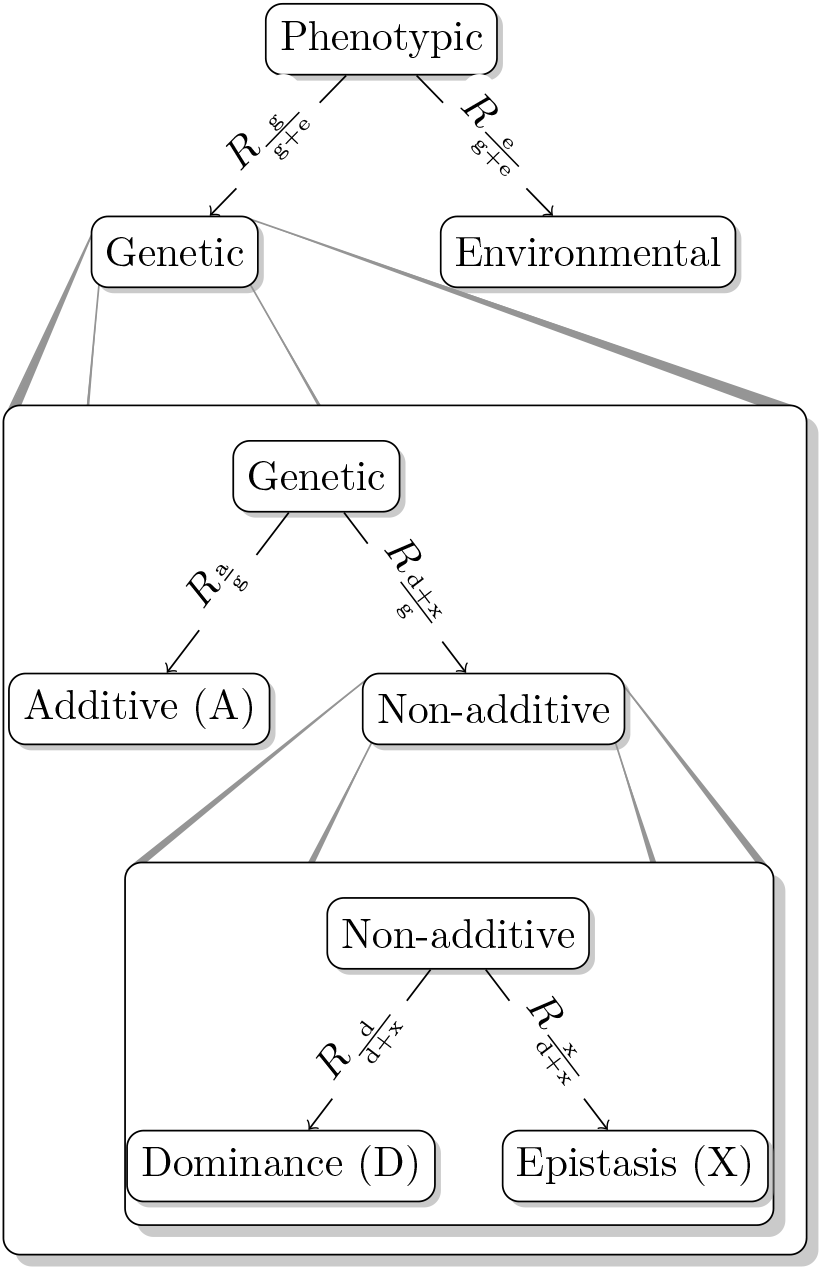
Tree structure visualizing the three possible model formulations A, AD and ADX. Edge labels illustrate where expert knowledge applies, namely on the relative magnitude of the genetic and environmental variation and the relative magnitude of the additive, dominance and epistasis variation.

The aim of this paper is to demonstrate the new approach and to evaluate the potential impact of using the approach along with expert knowledge in plant breeding. We first describe the genomic model and how to incorporate the expert knowledge in this model. To test the proposed approach, we first use a simulated wheat breeding program and evaluate inference stability, estimation of genetic values and variance components with different priors and with the standard maximum likelihood estimation. We also investigate the impact of dataset size. Then we apply the approach to a publicly available wheat yield dataset with 1,739 individuals from 11 different trials in 6 locations in Germany with varying amounts of observed phenotypes from Gowda *et al.* (2014) and Zhao *et al.* (2015). We use cross-validation to assess the accuracy of phenotype prediction when using the proposed priors in the model. A description of the simulated and real wheat breeding case studies, model fitting and analysis follows. Our key focus is to demonstrate how an analyst can take advantage of expert knowledge from literature or domain experts to enable robust genomic modelling of additive and non-additive variation. This focus involves specifying and visualizing the expert knowledge in an intuitive way. We then present the results and discuss the relevance of our work.

## Materials and Methods

### Genomic model

We model observed phenotypic values of *n* individuals ***y*** = (*y*_1_*, …, y_n_*) with the aim to estimate their genetic values and their additive and non-additive components. To this end we use the genomic information about the individuals contained in the observed single nucleotide polymorphism (SNP) matrix **Z**, where row *i* contains SNP marker genotypes for individual *i* coded additively with 0, 1, 2. We let **Z**_a_ be the column-centered **Z** where we have removed markers with low minor allele frequency, and let **Z**_d_ be the column-centered matrix obtained from **Z** after setting heterozygote genotypes to 1 and homozygote genotypes to 0.

We model the phenotypic observation *y*_*i*_ of individual *i* as

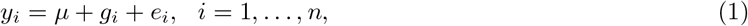

where *μ* is an intercept, *g*_*i*_ is the genetic value and *e*_*i*_ the environmental residual for individual *i*. We model the environmental residual as an independently and identically distributed Gaussian random variable, 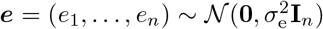, where 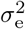 is the environmental variance and **I**_*n*_ is the *n* × *n* identity matrix. The intercept is typically well-identified from the data, and we specify the nearly translation-invariant prior 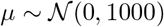.

We consider the simple additive model with *g*_*i*_ = *a*_*i*_ (*Model A*), and non-additive extension with dominance *g*_*i*_ = *a*_*i*_ % *d*_*i*_ (*Model AD*), and epistasis *g*_*i*_ = *a*_*i*_ + *d*_*i*_ + *x*_*i*_ (*Model ADX*). Here, ***a*** = (*a*_1_, …, *a*_*n*_), ***d*** = (*d*_1_, …, *d*_*n*_) and ***x*** = (*x*_1_, …, *x*_*n*_) respectively denote vectors of the additive, the dominance and the epistasis values for the individuals. Figure 1 shows the model structure for all three models, where every added component extends the model tree by one level. Moving from the root downwards, Model A is defined by the first split. Here only the additive value represents the genetic value. Model AD is defined by the first two splits, and as such has one level more. The genetic value splits into additive and non-additive values, where only the dominance value represents the non-additive value. Model ADX is defined by the complete tree and the non-additive value consists of both dominance and epistasis values.

We model the genetic values as 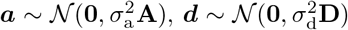 and 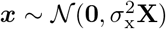, where 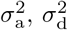 and 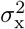 are the additive, dominance and epistasis variances, respectively. We specify the covariance matrices as 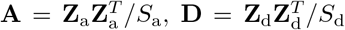 and **X**= **A** ☉ **A***/S*_x_ (we consider only additive-by-additive epistatis), where ☉ is the Hadamard product (Henderson, 1985; Horn, 1990; Gianola and de los Campos, 2008; Vitezica *et al.*, 2017). To incorporate our expert knowledge in a unified way, we scale the covariance matrices with *S*_a_, *S*_d_, and *S*_x_ according to Sørbye and Rue (2018). The idea of such scaling is not new, see Legarra (2016), Vitezica *et al.* (2017) and Fuglstad *et al.* (2020) for details. Finally, the phenotypic variance is 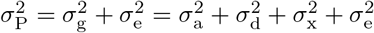.

### Expert knowledge about variance components

As highlighted in the introduction, there is prior information about the relative magnitude of the genetic and environmental variation and the relative magnitude of the additive, dominance and epistasis variation that can guide the construction of prior distributions. We specify this expert knowledge (EK) in a hierarchical manner:

**EK-pheno** informs on the split of phenotypic variation into genetic and enviromental variation. The proportion of genetic to phenotypic variation is denoted as 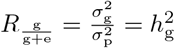, where 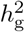 is the broad-sense heritability.
**EK-genetic** informs on the split of genetic variation into additive and non-additive variation. The proportion of additive to genetic variation is denoted as 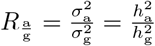, where 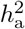 is the narrow-sense heritability.
**EK-nonadd** informs on the split of non-additive variation into dominance and epistasis variation. The proportion of dominance to non-additive variation is denoted as 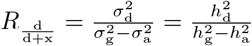, where 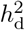 is the proportion of dominance to phenotypic variation.

Figure 1 illustrates where the respective expert knowledge in the form of relative magnitudes *R*_*_ applies. Of note, for Model A only EK-pheno is used, and EK-genetic is one 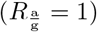 as non-additive effects are not considered in this model. Similarly, for Model AD only EK-pheno and EK-genetic are used as EK-nonadd is one 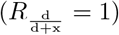.

Values for the relative magnitudes *R*_*_ will vary between study systems and traits in line with the expert knowledge. In this study our knowledge is based on the cited literature in the introduction and practical experience with the analysis of a range of datasets. We follow the fact that many complex traits in agriculture are under sizeable environmental effect and that additive effects capture most genetic variation by standard quantitative model construction. With this in mind we assume EK-pheno to be 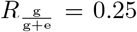, EK-genetic to be 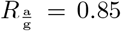 and EK-nonadd to be 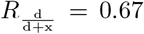. This implies 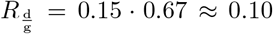 and 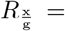 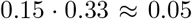. We emphasize that we use this information to construct prior distributions, i.e., these relative magnitudes are only taken as a guide and not as the exact magnitude of variance components. Fuglstad *et al.* (2020) show that the prior for the first partition, the broad-sense heritability 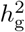, is not very influential.

We present two approaches for constructing a prior based on EK-pheno, EK-genetic and EK-nonadd: 1) a component-wise (comp) prior, that is placed independently on each variance parameter; and 2) a tree-based (tree) model-wise prior, that is defined jointly for all variance parameters. Both approaches are motivated by the concept of penalized complexity priors (Simpson *et al.*, 2017).

### Penalized complexity priors

A penalized complexity (PC) prior for a parameter *θ* is typically controlled by: 1) a preferred parameter value *θ*_0_ which is intuitive or leads to a simpler model; and 2) an idea on how strongly we believe in *θ*_0_. The PC prior shrinks towards *θ*_0_, unless the the data indicate otherwise. This is achieved by constructing the prior based on a set of well-defined principles, for details we refer to Simpson *et al.* (2017). PC priors can be applied to a standard deviation or variance, a proportion of variances, or other parameters such as correlations (Guo *et al.*, 2017).

The PC prior for a standard deviation (*σ*) of a random effect will shrink the standard deviation towards zero, that is, towards a simpler model without the corresponding random effect (assuming the prior mean of the effect is zero). This prior is denoted as 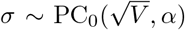 and results in an exponential distribution with rate parameter − 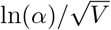. The subscript 0 in PC_0_(·) indicates that the prior shrinks towards *σ* = 0. To define the prior the analyst has to specify an upper value 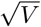 and a tail probability 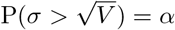 such that the upper-tail probability *α*. Here, we use *α* = 0.25 so the prior distribution is weakly-informative towards 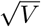, but shrinks to zero unless the data informs otherwise.

For a variance proportion *p* ∈ [0, 1] we denote the PC prior as *p* ∼ PC_0_(*R*). The numerical value *R* ∈ [0, 1] encodes the available expert knowledge about the proportion and is set as the median of the prior, i.e. P(*p* > *R*) = 0.5. The subscript 0 indicates that the prior shrinks towards *p* = 0. Shrinkage towards the median is achieved by the PC prior *p* ∼ PC_M_(*R*), where *R* has the same interpretation as for PC_0_(*R*). For PC_M_(*R*), we need to specify how concentrated the distribution is on logit-scale in the interval [logit(*R*) − 1, logit(*R*) + 1] around the median (see Fuglstad *et al.* (2020) for details). We allocated 75% probability to this interval.

The PC prior for a variance proportion depends on the structure of the two random components that are involved through their covariance matrices. We omit this in the notation for simplicity, and to emphasize that we chose to make the marginal priors on the proportions independent of each other. As the PC prior on proportions depends on the covariance matrix structure, it is application specific, and the priors do not correspond to common families of distributions such as the exponential or normal distributions (see Riebler *et al.* (2016); Fuglstad *et al.* (2020) for more details).

### Component-wise prior

In the component-wise setting we use a PC prior for each standard deviation parameter *σ*_*_. The PC prior on *σ*_*_ requires an upper value ⎷*V*_*_, so in addition to the relative magnitudes specified through EK-pheno, EK-genetic and EK-nonadd we need information on the magnitude of the phenotypic variance to set up the component-wise priors. For this purpose we could calculate the empirical phenotypic variance *V*_P_ from a separate dataset, which is a trial study or a study believed to exhibit similar phenotypic variance as the study at hand. From this we can define the upper values for the individual PC priors. For example, to formulate priors for Model A we use EK-pheno to find 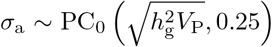 and 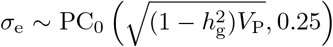. For Model AD we need EK-pheno and EK-genetic to formulate the priors, and for Model ADX, the most complex model, we take into account all available expert knowledge.

We follow the tree-structure shown in Figure 1 downwards to define the upper values, and multiply the relative magnitudes on the edges leading to the respective leaf nodes. For Model ADX this leads us to:

- 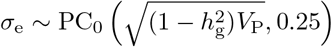,
- 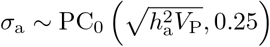,
- 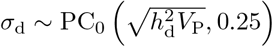, and
- 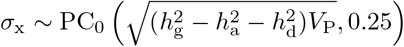.

Combining the available expert knowledge procedure with the three different genomic models gave us settings we denote as A-comp*, AD-comp* and ADX-comp*. We have contrasted these settings with a default component-wise PC prior proposed by Simpson *et al.* (2017) with 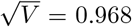 and *α* = 0.01 on all variance parameters, which gave us settings denoted as A-comp, AD-comp and ADX-comp. Preliminary analyses showed that the inferences for AD-comp, AD-comp*, ADX-comp and ADX-comp* are not stable, the methods are not robust in the sense that they did not avoid estimating spurious non-additive effects, and we do not present results from these settings. The priors for A-comp* and A-comp are plotted in Figure 2 using 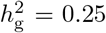 and *V*_P_ = 1. If *V*_P_ takes another value, we simply rescale the *x*-and *y*-axes; the shape of the prior stays the same. In the simulated case study, we will use *V*_P_ = 1.86. The priors are equal on all standard deviations for A-comp, AD-comp and ADX-comp. The priors for AD-comp* and ADX-comp* can be seen in Figures S1 and S2 in File 1 in the Supplemental materials. See Note S1 in File S1 for a detailed description of the component-wise prior and posterior distributions for Model A and Model AD.

**Figure 2:**
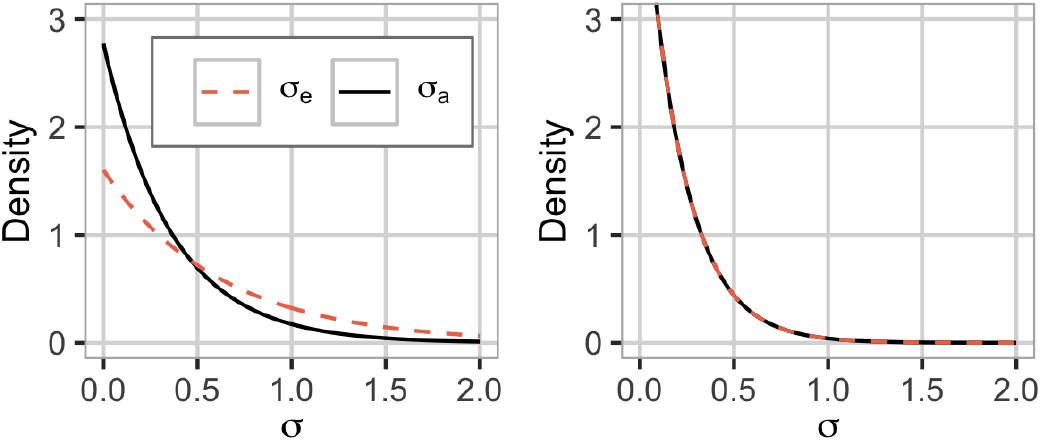
The prior used for the A-comp*^*a*^ (left) and A-comp^*b*^ (right) settings. For A-comp*, we use 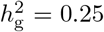. We have plotted the priors for *V*_P_ = 1. For A-comp, additive and environmental variances have the same prior.

### Tree-based model-wise prior

In the model-wise setting, we shift the focus in Figure 1 from the leaf nodes to the splits. In other words, a shift from the component-wise perspective of variances associated with different sources of variation to a model-wise perspective of how these variances arise as a hierarchical decomposition of the phenotypic variance. This provides a complementary way to construct priors where EK-pheno, EK-genetic and EK-nonadd are directly incorporated at the appropriate levels in the tree structure. We achieve this by applying the hierarchical decomposition (HD) prior framework of Fuglstad *et al.* (2020). We focus the presentation on the essential ideas for understanding and successfully applying the priors, and provide the comprehensive and mathematical description in Note S1 in File S1. We emphasize that in the following *p*_*_ denotes an actual variance proportion that we will infer (along with variances), while *R*_*_ denotes expert knowledge for this proportion.

We first assign a marginal prior for the decomposition of variances in the lowest split, and then move step-wise up the tree assigning a prior to the decomposition of variance in each split conditional on the splits below it. The bottom-up process ends with the assignment of a prior to the root split, and the result is a joint prior for the decomposition of phenotypic variance into the different sources of variance. In the final step, we assign a prior for phenotypic variance 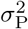 that is conditionally independent of the prior on the decomposition of the phenotypic variance.

We follow Fuglstad *et al.* (2020) and simplify the prior at each split by conditioning on expert knowledge from the lower splits. For example, the prior for 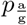 a is constructed under the assumption that 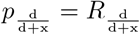; that is, 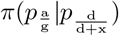 is replaced with 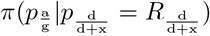. Note that even though we construct the prior from the bottom and up, the arrows in the tree indicate how the phenotypic variance is distributed in the model from the top down. This means that the amount of, for example, dominance variance 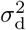 depends on the variance partitions further up, since 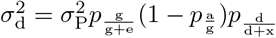 following the tree structure (Figure 1).

In this study, we assumed that at the lower levels the model shrinks towards the expert knowledge EK-nonadd and EK-genetic by using PC_M_(·) priors. Further, at the top level we use a PC_0_(·) prior to shrink towards the environmental effect unless the data indicates otherwise to reduce overfitting. Note that we could have chosen different assumptions. To obtain a prior fulfilling our assumptions, we follow four steps:

1. we use a PC_M_(·) prior for the proportion of dominance to non-additive variance with median 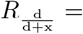 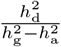 (EK-nonadd),
2. we use a PC_M_(·) prior for the proportion of additive to genetic variance with median 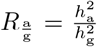 (EK-genetic),
3. we use a PC_0_(·) prior for the proportion of genetic to phenotypic variance with median 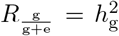 (EK-pheno), and
4. we achieve scale-independence through the non-informative and scale-invariant Jeffreys’ prior for the phenotypic variance 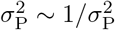.

This construction gives the joint prior 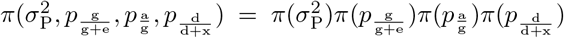 for Model ADX, where the conditioning on expert knowledge from lower splits is omitted to simplify notation. We denote this setting as ADX-tree* and show this prior in Figure 3 for 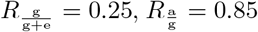 and 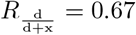. Note that the model-wise priors with expert knowledge are dependent on the covariance matrices of the modelled effects and are therefore dataset specific (Fuglstad *et al.*, 2020), and the plots of these priors thus pertain to one specific dataset. The spike at *p*=1 for 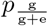 in Figure 3 is an artifact of the parameterization chosen for visualization and does not induce overfitting; see Fuglstad *et al.* (2020) for details. See Note S1 in File S1 for a detailed description of the model-wise prior and posterior distributions for Model A and Model AD.

**Figure 3:**
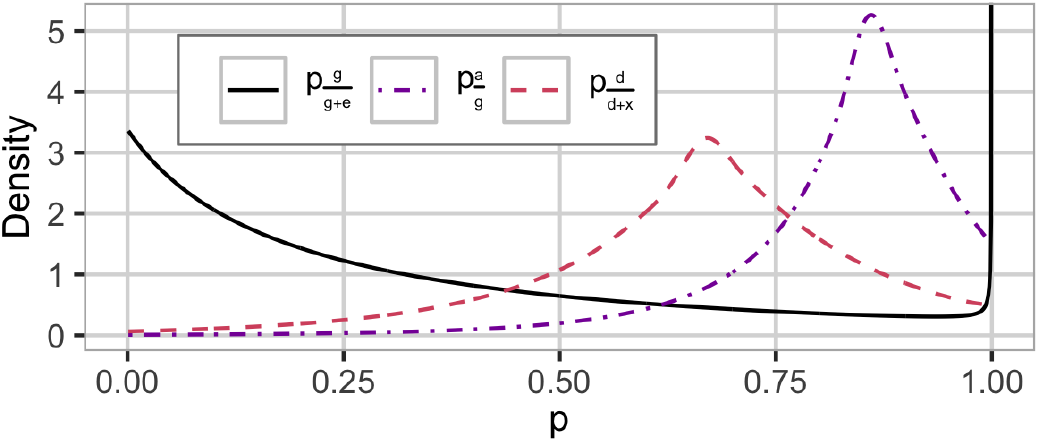
The HD prior used in the ADX-tree* ^*a*^ setting with the proportion of genetic to phenotypic variance 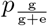, additive to genetic variance 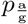 and dominance to non-additive variance 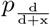. We use 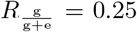, 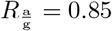 and 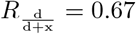. This is a dataset specific prior.

We explored the influence of alternative expert knowledge. In addition to the previously stated values for EK-pheno, EK-genetic and EK-nonadd we also tested 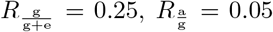, and 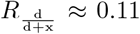 (so 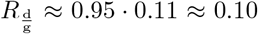 and 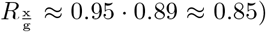. The constructions follows the description above but with these relative magnitudes instead. We denote this setting as ADX-tree-opp*, as it expresses almost opposite or “wrong” beliefs compared to ADX-tree* setting, and show the prior in Figure S3 in File S1 in the Supplemental materials.

For Model AD the non-additive effect only consists of dominance, and the variance is attributed to the different effects as visualized by the top and middle split in Figure 1. We construct a prior using EK-pheno and EK-genetic with 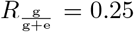 and 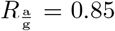 and denote this setting AD-tree*. The prior is shown in Figure 4.

**Figure 4:**
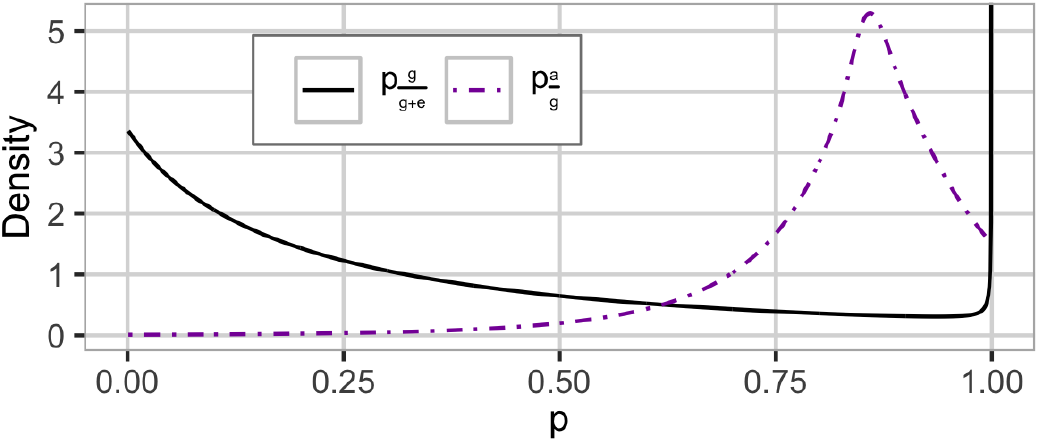
The HD prior used in the AD-tree* ^*a*^ setting with the proportion of genetic to phenotypic variance 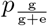 and additive to genetic variance 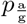. We use 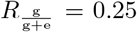 and 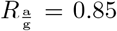. This is a dataset specific.

For Model A the genetic variance is not decomposed to different sources and the distribution of the phenotypic variance can be visualized using the top split in Figure 1. We use EK-pheno with 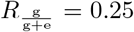 to construct a prior for the proportion of genetic to phenotypic variance and denote this setting as A-tree*. We show this prior in Figure 5.

**Figure 5:**
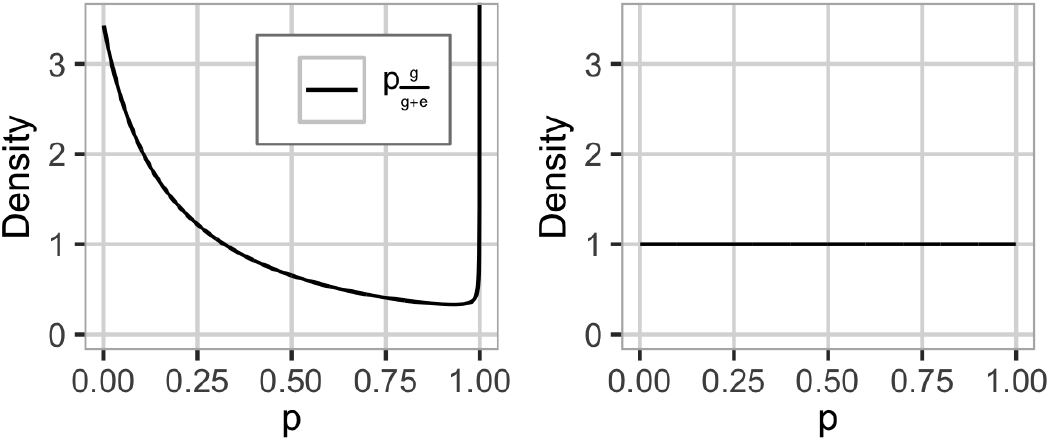
The prior for the proportion of genetic to phenotypic variance 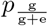 for the A-tree*^*a*^ (left) and ^*b*^ (right) settings. We use 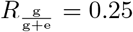. A-tree* is a dataset specific prior.

We compared the model-wise prior with expert knowledge to a default prior with no expert knowledge by constructing an HD prior using the exchangeable Dirichlet prior on the proportion of phenotypic variance attributed to each of the sources of variance following Fuglstad *et al.* (2020). For Model A we use a uniform prior, which is a special case of the symmetric Dirichlet(*m*) prior with *m* = 2, on the proportion of genetic to phenotypic variance 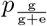 and denote this setting as A-tree (see Figure 5). For Models AD and ADX we use Dirichlet(3) and Dirichlet(4) priors on the proportions, respectively, and denote these settings AD-tree and ADX-tree. These priors do not induce shrinkageards any of the effects, and assume that each model and effect contributes equally to the phenotypic variance, which due to the lack of expert knowledge did not lead to stable inference for Models AD and ADX. We do not show results from these settings. The tree structure and prior for AD-tree and ADX-tree can be seen in Figures S4 and S5 in File S1, respectively. We summarize the model-wise priors that will be used in the following in Table 1.

**Table 1:** Summary of the model-wise (tree-based) prior distributions on proportionsup^*a,b*^ and total phenotypic variance.

| **Additive**<br>**(Model A, $g_i = a_i$)** |  | **Additive and dominance**<br>**(Model AD, $g_i = a_i + d_i$)** | **Additive and non-additive**<br>**(Model ADX, $g_i = a_i + d_i + x_i$)** |
| --- | --- | --- | --- |
| *Default* | *Expert* | *Expert* | *Expert* |
| A-tree:<br>$p_{\frac{g}{g+e}} \sim \text{Dirichlet}(2)$<br>$\sigma_P^2 \sim \text{Jeffreys}'$ | A-tree*:<br>$p_{\frac{g}{g+e}} \sim \text{PC}_0\left(R_{\frac{g}{g+e}}\right)$<br>$\sigma_P^2 \sim \text{Jeffreys}'$ | AD-tree*:<br>$p_{\frac{g}{g+e}} \sim \text{PC}_0\left(R_{\frac{g}{g+e}}\right)$<br>$p_{\frac{a}{g}} \sim \text{PC}_M\left(R_{\frac{a}{g}}\right)$<br>$\sigma_P^2 \sim \text{Jeffreys}'$ | ADX-tree*:<br>$p_{\frac{g}{g+e}} \sim \text{PC}_0\left(R_{\frac{g}{g+e}}\right)$<br>$p_{\frac{a}{g}} \sim \text{PC}_M\left(R_{\frac{a}{g}}\right)$<br>$p_{\frac{d}{d+x}} \sim \text{PC}_M\left(R_{\frac{d}{d+x}}\right)$<br>$\sigma_P^2 \sim \text{Jeffreys}'$ |

<sup>a</sup> $p \sim \text{PC}_0(R)$ describes a PC prior for a variance proportion that has median equal to $R$ and a preference for the variance proportion being equal to 0.
<sup>b</sup> $p \sim \text{PC}_M(R)$ describes a PC prior for a variance proportion with median $R$ and a preference for the variance proportion being equal to the median $R$ , with 75% probability in $[\text{logit}(R) - 1, \text{logit}(R) + 1]$ around the median on logit-scale.

### Simulated case study

We applied the described genomic model (1) with the above mentioned priors to a simulated case study that mimics a wheat breeding program to investigate the properties of the different settings. We simulated the breeding program using the R package AlphaSimR (Faux *et al.*, 2016; Gaynor, 2019) and closely followed the breeding program descriptions of Gaynor *et al.* (2017) (see their Figure 3) and Selle *et al.* (2019).

Specifically, we simulated a wheat-like genome with 21 chromosomes, selected at random, 1, 000 SNP markers and 1, 000 causal loci from each chromosome and used this genome to initiate a breeding program with breeding individuals. Every year we have used 50 fully inbred parents to initiate a new breeding cycle with 100 random crosses. In each cross we have generated 10 progenies and selfed them to generate 1, 000 F2 (second filial) individuals, which were selfed again to generate 10, 000 F3 (third filial) individuals. We reduced the 10, 000 F3 individuals in four successive selection stages (headrow, preliminary yield trial, advanced yield trial and elite yield trial) with 10% selection intensity in each stage. In the headrow stage, we simulated a visual selection on a phenotype with the heritability of 0.03. For the preliminary, advanced and elite yield trials we respectively simulated selection on phenotype with heritability 0.25, 0.45 and 0.62. We used the 50 individuals with the highest phenotype values from the last three selection stages as parents for the next breeding cycle.

We ran the simulation for 30 years. At year 1, we set the following variances for the population of the 50 parents: additive variance of 1.0, dominance variance of 0.5, and epistasis variance of 0.1. We set the environmental variance to 6.0 for all stages and years. We ran the simulation for 20 years as a “burn-in” to obtain realistic breeding data under selection. We then ran the simulation for another 10 years with selection on phenotype. We removed the SNP markers with minor allele frequency below 5%. We did not use the models for selection.

### Real case study

We also applied the described genomic model (1) to the publicly available Central European wheat grain yield data from Gowda *et al.* (2014) and Zhao *et al.* (2015). In short, the data consists of 120 female and 15 male parent lines, which were crossed to create 1,604 hybrids. The parents and hybrids were phenotyped for grain yield in 11 different trials in 6 locations in Germany. The number of observed phenotypes for the parents and hybrids vary between the trials, i.e., some datasets have more observed phenotypes than others, ranging from 834 to 1,739 (see Table S1 in File S1 in the Supplemental materials). The parents and hybrids have genotype data for 17,372 high-quality SNP markers.

In the real case study we analyzed the performance of the tree-based priors using expert knowledge (tree*) for the additive model (A), the additive and dominance model (AD), and the additive and non-additive model (ADX). We used the same as in the simulation study: 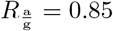 and 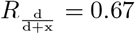. We have however used a higher value in EK-pheno, 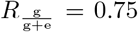, in line with *Reif et al.* (2011) - later stage trials tend to have higher heritablity than early stage trials. Again, we emphasize that these values are only used to construct prior distributions and are not taken as literal proportions. The resulting priors can be seen in Figure S6 in File S1.

### Implementation details

We fitted the models with a Bayesian approach through the R package RStan (Carpenter *et al.*, 2017; Stan Development Team, 2019). This package provides a sampling algorithm that uses the No-U-Turn sampler, a variant of Hamiltonian Monte Carlo, and only requires that the user specifies the joint posterior distribution up to proportionality, without having to write a sampling algorithm. See Note S1 in File S1 in the Supplemental materials for details. Sampling methods such as Markov Chain Monte Carlo and Hamiltonian Monte Carlo have guaranteed asymptotic accuracy as the number of drawn samples go to infinity. However, in an applied context with finite computational resources, it is hard to assess this accuracy. Betancourt (2016) developed a diagnostic metric for the Hamiltonian Monte Carlo, called divergence, that indicates whether the sampler is able to transition through the posterior space effectively or not, where in the latter case the results might be biased (we show an example on this in the results).

We also fitted Models A, AD and ADX with the maximum likelihood (ML) approach using the low-storage BFGS (Broyden–Fletcher–Goldfarb–Shanno) algorithm through the R package nloptr (Nocedal, 1980; Liu and Nocedal, 1989; Johnson, 2020). This approach does not use priors. We denote them as A-ML, AD-ML and ADX-ML and use them as a base-line for comparison because maximum likelihood is a common approach in the literature. Inference for ADX-ML was not robust, and we do not present results from this setting. At last, we compared the model results to the performance of selection based solely on phenotype where we treat the phenotype as a point estimate of the genetic value.

### Performance assessment

For the simulated case study, we ran the breeding program simulation 4,000 times and fitted the model and prior settings in each of the last 10 years of simulation (40,000 model fits in total) at the third selection stage (advanced yield trial) in the program. Here we had 100 individuals each with five replicates and the goal was to select the 10 genetically best individuals for the fourth, last, stage. For each model fit we evaluated: (i) robustness of method, (ii) the accuracy of selecting the genetically best individuals, (iii) the accuracy of estimating the different genetic values and (iv) the accuracy of estimating the variance parameters. We evaluated the fits against the true (simulated) values.

We measure how *robust* the method (model and inference approach) is, i.e., to which degree it avoids estimating spurious non-additive effects, in *stability of inference*. For the stability of inference of the Bayesian approach with Stan we used the proportion of analyses that had stable inference (which we define as at least 99% samples where no divergent transitions were observed) for each model and prior setting. For the stability of inference of the maximum likelihood approach we used the proportion of analyses where the optimizer algorithm converged.

For the accuracy of selecting the genetically best individuals we ranked the best 10 individuals based on the estimated genetic value or estimated additive value, and counted how many were among the true genetically best 10 individuals based on the true genetic value or true additive value. We used the posterior mean of the effects as estimated values for ranking. Selection on the genetic value indicated selection of new varieties, while the selection on the additive value indicated selection of new parents.

For the accuracy of estimating the different genetic values (total genetic, additive, dominance and epistasis values) we used Pearson correlation and continuous rank probability score (CRPS, Gneiting and Raftery, 2007). With the correlation we measured how well posterior means of genetic values correlated with true values (high value is desired). This metric works with point estimates and ignores uncertainty of inferred posterior distributions of each individual genetic value. The CRPS is a proper scoring rule and as such measures a combination of bias and sharpness of the posterior distribution compared to true values (low value is desired). Specifically, CRPS integrates squared difference between the cumulative estimated posterior distribution and the true value over the whole posterior distribution (Gneiting and Raftery, 2007). See Selle *et al.* (2019) for a detailed explanation of CRPS used in a breeding context. In the case of phenotypic selection, we have a phenotype value for selection candidates, which is a point estimate of the genetic value, and its CRPS then reduces to the mean absolute error between the true genetic values and the phenotype. The accuracy of the estimates of the variance parameters was assessed by dividing them by the true genetic variances for each of the 10 years from the simulated breeding program (a value close to 1 is desired). This is not done for phenotype selection.

To test the effect of dataset size on inference, we ran the breeding program an additional 1,000 times and fitted the models to *n* = 700, 600, …, 100 individuals in the preliminary stage (instead to 100 individuals in the advanced stage) at year 21. We used the settings with tree-based expert knowledge priors and the maximum likelihood approach and investigated the performance of the methods for increasing number of observations by evaluating the robustness, and the accuracy of estimating the different genetic values and variance parameters.

We analyzed the real case study with the same models and tree-based expert knowledge priors and focused on the ability of predicting observed phenotypes in a cross-validation shceme. We performed 5-fold cross-validations five times for each of the 11 trials independently. For each fold in each cross-validation, we predicted the held-out phenotypes (their posterior distribution involving intercept, genetic value and environmental variation), and calculated the correlation between the point predictions and the observed phenotypes, and the CRPS using the phenotype posterior prediction distributions and the observed phenotypes available for each trial. We note that phenotype posterior predictions involve environmental variation, which does not affect point predictions and correlations, but affects the CRPS as the whole distribution of the prediction is involved in the calculations. We also looked at the posterior medians of the model variances. Of note, in contrast to the simulated case study the genetic effects are unknown for real data, so that we cannot assess the estimation accuracy of the effects.

### Data and code availability

We provide code to simulate the data described in the simulated case study (File S2 in the Supplemental materials). We also provide example code to fit the models presented in this paper together with an example dataset (File S3). In the real case study we used data from Gowda *et al.* (2014) (SNP genotypes) and Zhao *et al.* (2015) (phenotypes), and provide code for fitting the models in File S4, including the folds used in the cross-validation. The Supplemental materials are available at figshare: https://doi.org/10.6084/m9.figshare.12040716.

## Results

### Simulated case study

In the simulated case study the model-wise priors and expert knowledge improved the inference stability of the non-additive models and the selection of the genetically best individuals, but did not significantly improve the accuracy of estimating different genetic values for all individuals or for variance components.

#### Robustness and stability

Table 2 shows the proportion of stable model fits by model and prior setting. The model-wise priors combined with expert knowledge improved the inference stability of the additive and dominant (AD) model and the non-additive (ADX) model to the level of stability of the additive (A) model and phenotypic selection. Phenotypic selection does not depend on a model fit to a dataset and therefore had the highest method robustness by definition. This maximum level of robustness was matched by the simple additive model with the model-wise prior with and without using expert knowledge (A-tree* and A-tree) and with the standard maximum likelihood approach (A-ML). This high model robustness was followed closely by fitting the more complicated non-additive and additive and dominance models with model-wise prior and expert knowledge (ADX-tree* and AD-tree*). The Bayesian approach using component-wise priors with expert knowledge (A-comp*), the additive and dominance model with the maximum likelihood approach (AD-ML), the component-wise priors without expert knowledge (A-comp), and the model-wise prior with wrong/opposite expert knowledge (ADX-tree-opp*) also resulted in satisfactory robustness, but then the proportion of model fits with stable inference started to decrease. The robustness of the additive and dominance model and the non-additive model with default component-wise priors (AD-comp and ADX-comp) was improved by using the model-wise priors (AD-tree and ADX-tree), and even further by expert knowledge (AD-comp* and ADX-comp*), but neither they nor the non-additive model fitted with maximum likelihood (ADX-ML) had more than 80% stable model fits.

**Table 2:** Method robustness measured in stability of inference ^*a*^ by model and prior setting.

| Setting (abbreviation) | Stability |
| --- | --- |
| Phenotype selection | 1.00 |
| Add. tree expert (A-tree*) | 1.00 |
| Add. tree default (A-tree) | 0.99 |
| Add. maximum likelihood (A-ML) | 0.99 |
| Non-add. tree expert (ADX-tree*) | 0.98 |
| Add. + dom. tree expert (AD-tree*) | 0.97 |
| Add. comp. expert (A-comp*) | 0.94 |
| Add. + dom. maximum likelihood (AD-ML) | 0.88 |
| Add. comp. default (A-comp) | 0.86 |
| Non-add. tree expert opposite (ADX-tree-opp*) | 0.86 |
| Non-add. comp. expert (ADX-comp*) | 0.80 |
| Non-add. maximum likelihood (ADX-ML) | 0.79 |
| Add. + dom. comp. expert (AD-comp*) | 0.69 |
| Add. + dom. tree default (AD-tree) | 0.51 |
| Non-add. tree default (ADX-tree) | 0.23 |
| Add. + dom. comp. default (AD-comp) | 0.13 |
| Non-add. comp. default (ADX-comp) | 0.04 |

<sup>a</sup>as a proportion of analyses with less than 1% divergences for the Bayesian approach and as a proportion of analyses with convergence for the maximum likelihood approach.

The reason for deteriorated robustness of some model and prior settings is that genetic (especially the non-additive) and environmental effects can be partially confounded, which limits the exploration of the posterior when using the Bayesian approach or limits convergence of mode-seeking algorithms when using the maximum likelihood approach. We show the partial confounding with images of the covariance matrices for additive, dominance, epistasis and environmental sources of variation for one dataset in Figure S7 in File S1 in the Supplemental materials. Figure S8 shows joint posterior samples for the epistasis and environmental variance for model ADX with model-wise priors with and without expert knowledge (ADX-tree* and ADX-tree) for one dataset. Without a robust method (this includes both the model and inference approach), the posterior distribution becomes difficult to explore, and this is also supported by the divergence diagnostics (Table 2). The posterior of the ADX-tree setting is bimodal and the sampler has not been able to sample with convergence due to confounding.

We do not present results from the settings with 80% or less stable model fits (see Table 2) in the following. Note that Table 2 includes all model abbreviations used. For each setting, the breeding programs that did not result in stable inference were removed from the results.

#### Selecting best individuals

Figure 6 shows the accuracy of selecting individuals with the highest genetic value (variety selection, Figure 6a) and with the highest additive value (parent selection, Figure 6b). The model-wise priors exploiting expert knowledge improved the selection of the genetically best individuals significantly, and the model choice was important for different breeding aims. The settings with the additive and dominance model and the non-additive model with model-wise expert knowledge (AD-tree* and ADX-tree*) performed significantly better in selection of new varieties than the others, which followed in order from A-tree, A-tree*, A-comp*, A-comp, A-ML, ADX-tree-opp* and AD-ML (see Table 2 for abbreviations). The differences between the settings were small, but they all performed significantly better than sole phenotype selection, which is sensitive to environmental noise. For the selection of new parents the simpler additive model performed the best, and the model-wise priors improved its performance (A-tree, A-tree* and A-comp*). Wrong expert knowledge harmed the parent selection (ADX-tree-opp*), but it still outperformed sole phenotype selection.

**Figure 6:**
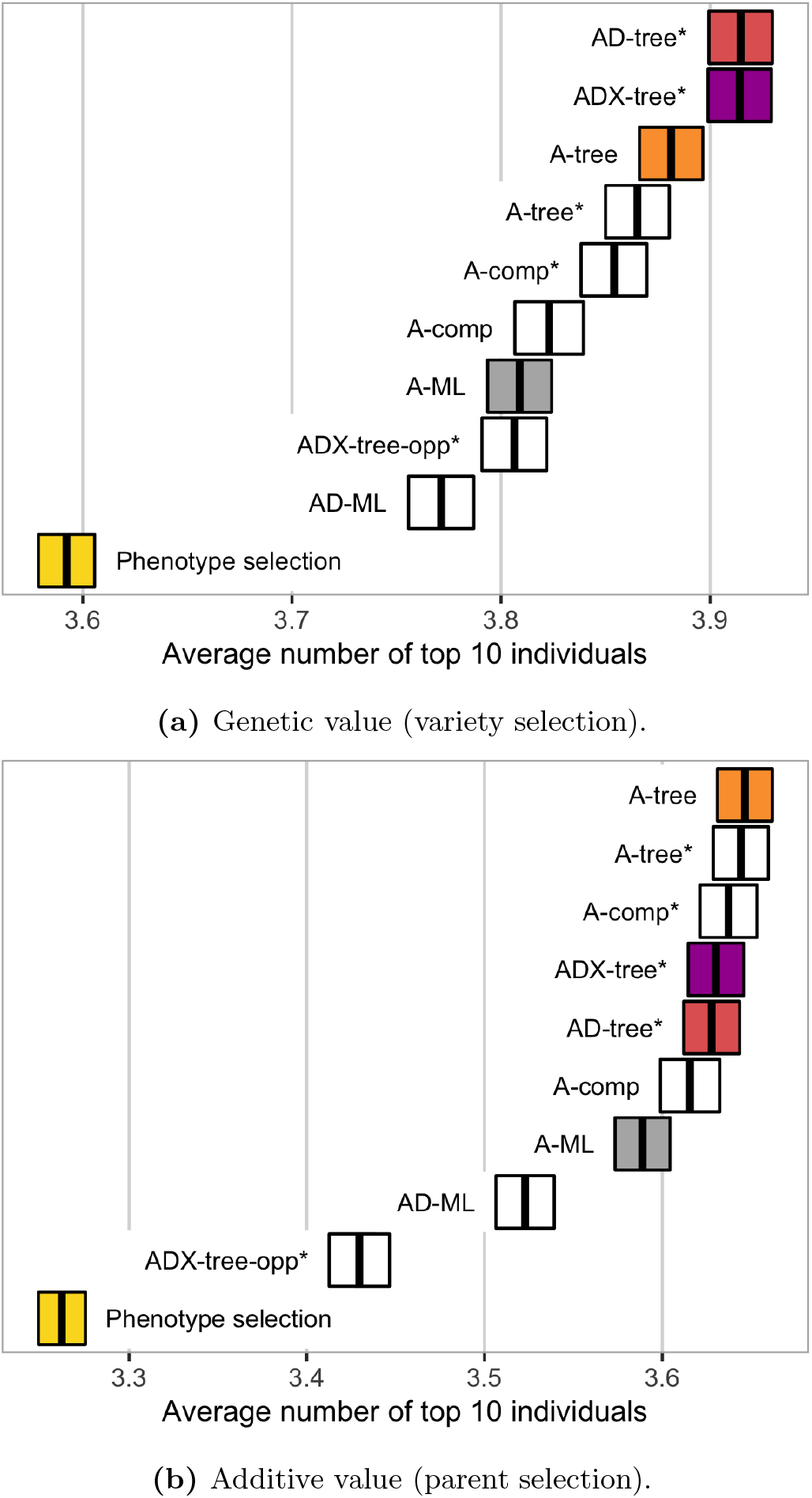
Accuracy of selecting individuals with the highest (a) genetic value (for variety selection) and (b) additive value (for parent selection) by model and prior setting - measured with the number of the top 10 true best individuals among the top 10 selected individuals (average ± two standard errors over replicates).

#### Estimation

We summarize the remaining results here, and include a detailed description of the results for the additive model with model-wise default prior (A-tree) and the maximum likelihood approach (A-ML), the additive and dominance model and the non-additive model with model-wise expert knowledge prior (AD-tree* and ADX-tree*), in addition to phenotype selection, in Note S2, and provide the complete results for all settings in Figures S9-S14 in File S1 in the Supplemental materials.

While using the model-wise priors and expert knowledge significantly improved the selection of the genetically best individuals compared to the maximum-likelihood approach, it did not significantly improve the accuracy of estimating different genetic values across all individuals (Figures S9 and S10). There was a tendency for the Bayesian models to perform better than the models fitted with the maximum likelihood approach, but the variation between replicates was larger than than the variation between the settings. All models outperformed phenotype selection, where we treat the phenotype as a point estimate of the genetic value.

Figure S11 shows that the variance component estimates varied considerably around the true values for all models and prior settings. The estimates from the Bayesian inference showed slightly larger biases and smaller variances than maximum likelihood estimates. Estimates for epistasis variance were considerably more underestimated than for the dominance variance.

The inference stability did not increase with increasing number of observations for any of the models fitted with the maximum likelihood approach. The Bayesian models with model-wise expert knowledge priors had the same high inference stability as in Table 2. The variation between replicates decreased for the variance estimates (Figure S12) and the correlation and continuous rank probability score (CRPS) improved for all models for increasing number of observations (Figures S13 and S14). 700 observations was not enough for the maximum likelihood approach to obtain a bias in dominance and epistasis variance estimates as low as the Bayesian approach (Figure S15), indicating that the need for good prior distributions is still there, but decreases with increasing number of observations.

### Real case study

The Bayesian approach with model-wise expert knowledge priors performed at least as good as or better than the maximum likelihood (ML) approach. Figure 7 shows the predictive ability of phenotypes measured with correlation and CRPS from three trials in Seligenstadt (Sel13 and Sel12) and Hadmersleben (Had12) over the five 5-fold cross-validations. These trials had phenotype observations for 1,739 (Sel13), 834 (Sel12) and 1,738 (Had12) parents and hybrids. We include correlation and CRPS for all 11 trials in Figures S15 and S16 in File S1 in the Supplemental materials. The maximum likelihood approach was as good as the Bayesian approach in the Sel13 trial where all phenotypes were observed for the parents and hybrids, but in the Sel12 trial, which consists of only 834 out of 1,739 observed phenotypes, the maximum likelihood approach had worse predictive ability for the additive model (A), and slightly worse for the non-additive model (ADX). In the Had12 trial with practically no unobserved phenotypes, the maximum likelihood approach is outperformed by the Bayesian approach for the non-additive model due to overfitting through overestimation of the epistasis variance (see Figure 8). The results from the additive and dominance (AD) model did not differ from the results from the additive and non-additive model, and we to not discuss them here, but include the results from AD-tree* and AD-ML in File S1 (Figures S15-S17).

**Figure 7:**
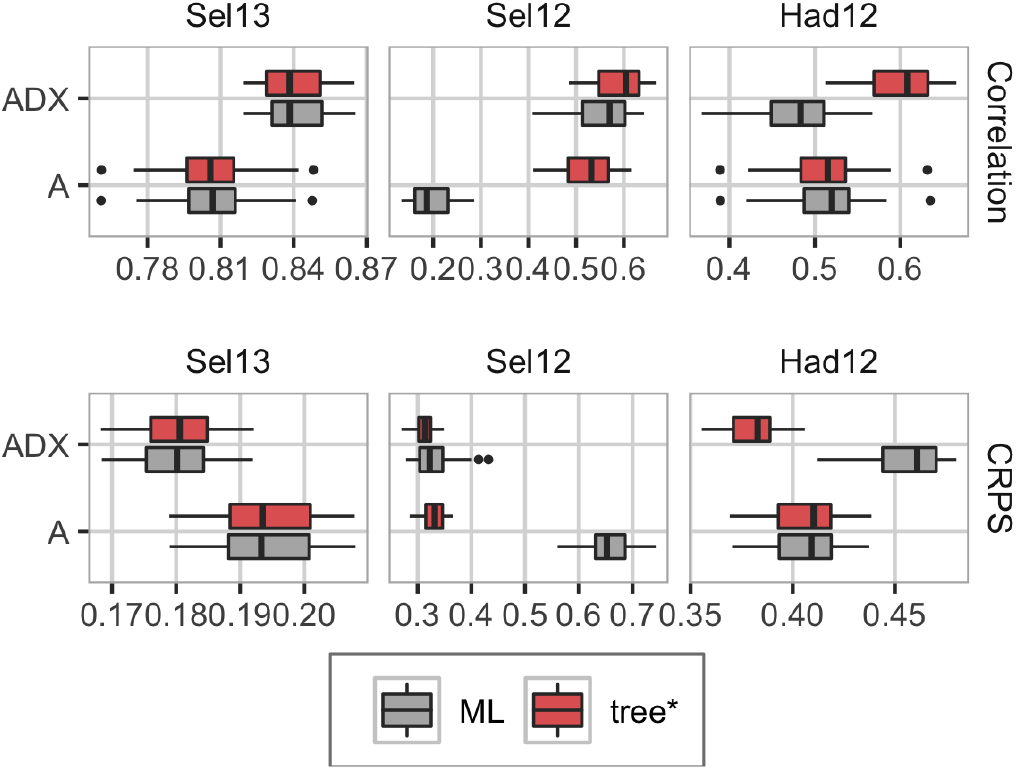
Phenotype prediction ability measured with correlation (top; high value desired), and continuous rank probability score (bottom; low value desired) from three of the trials in the real case study (boxplots show variation over the cross-validations and folds). Left: Sel13 (1,739 observations), middle: Sel12 (834 observations), right: Had12 (1,738 observations).

**Figure 8:**
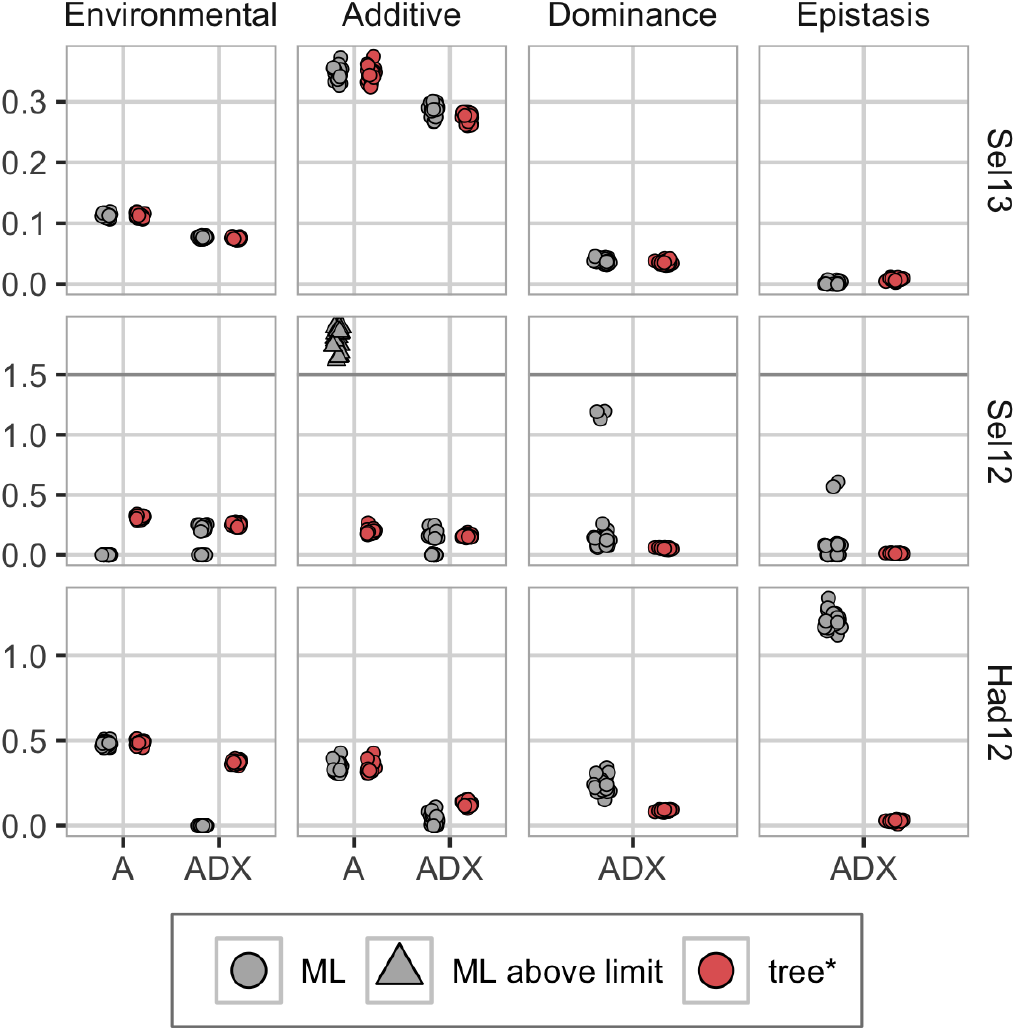
Posterior median variances from the real case study for three of the trials for the five 5-fold cross-validations. Top: Sel13 (1,739 observations), middle: Sel12 (834 observations), bottom: Had12 (1,738 observations). For Sel12, the A-ML is overestimating the additive variance so badly (values over 400) that we have truncated the y-axes at 1.5 to highlight the other results.

We explored reasons for the bad performance of A-ML in the Sel12 trial. The maximum likelihood optimizer returned a converge error message for two of the total 25 folds (we removed these model fits from all the results). However, the severe overestimation of the additive variance shown in Figure 8 indicates that the optimizer did not find the global maximum, but rather a local one. A closer investigation of the variance estimates showed that the optimizer got “stuck” at the lower boundary values (20 for the environmental and 50 for the other variances on a logarithmic scale). We gave 0 as initial value for the intercept and log-variances for both the Bayesian and maximum likelihood approach, however, the latter did not converge.

In Figure 8 we see that for Sel13 the approaches are in agreement on the variance estimates. With a dataset with many unobserved phenotypes (Sel12), the additive model fitted with the maximum likelihood approach (A-ML) estimated the environmental log-variance at −20, and in compensation severely overestimated the additive variance. The non-additive model fitted with maximum likelihood (ADX-ML) had the same underestimation of the environmental variance for some folds, but compensated with non-additive effects. This indicates overfitting and means that predictions from such are based solely on genetic values, and no environmental effects, which gives misleading predictions. ADX-ML was also underestimating the environmental variance for the data from Had12, and compensated this variance with the dominance and epistasis effects. We reran the maximum likelihood optimizer with initial values set to posterior medians from the corresponding Bayesian models. In this case, the maximum likelihood approach was not outperformed by the Bayesian approach (see Figures S15 and S16 in File S1). The variance estimates for all environments can be seen in Figure S18.

In Figures S15 and S16 we see that the trend is the same for most of the trials; for datasets where we have observed most of the phenotypes for the parents and hybrids, the maximum likelihood and Bayesian approaches are in general performing equally, and we gain predictive accuracy by including non-additive effects, but as soon as there are many unobserved phenotypes, such as for Boh12 and Sel12 (see Table S1 for information about all trials), the maximum likelihood approach is deteriorating. For the Had12, Had13 and Hhof13 trials, which has few unobserved phenotypes but still has poor predictive abilities for the non-additive model (ADX), the maximum likelihood approach has problems with overfitting (see Figure S17). The model underestimates the environmental variance and attributes this variation to the dominance and epistasis effects.

## Discussion

In this study we have introduced new priors for robust genomic modelling of additive and non-additive variation based on the penalized complexity prior (Simpson *et al.*, 2017) and hierarchical decomposition prior (Fuglstad *et al.*, 2020) frameworks. In the simulated case study, the new priors enabled straightforward use of expert knowledge, which in turn improved the robustness of genomic modelling and the selection of the genetically best individuals in a wheat breeding program. However, it did not improve the overall accuracy of estimating genetic values for all individuals or for variance components. In the real case study, the new priors improved the prediction ability, especially for trials with fewer observations, and they reduced overfitting. These results highlight three points for discussion: (i) expert-knowledge priors for genomic modelling and prediction, (ii) the importance of priors for breeding and (iii) limitations of our work.

### Expert-knowledge priors for genomic modelling and prediction

Genomic modelling is challenging due to inherent high-dimensionality and pervasive correlations between loci and therefore requires substantial amounts of information for robust estimation. Most genomes harbour millions of segregating loci that are highly or mildly correlated. While estimating additive effects at these loci is a challenging task in itself (e.g., Visscher *et al.*, 2017; Young, 2019), estimating dominance and epistasis effects is an even greater challenge (e.g., Misztal, 1997; Zhu *et al.*, 2015; de los Campos *et al.*, 2019). One challenge in estimating the interactive dominance and epistasis effects is that they are correlated with the main additive effects and all these effects are further correlated across nearby loci (Mäki-Tanila and Hill, 2014; Hill and Mäki-Tanila, 2015; Vitezica *et al.*, 2017). Information to estimate all these locus effects and corresponding individual values has to inherently come from the data, but could also come in a limited extent from the expert knowledge. There is a wealth of expert knowledge in genetics (e.g., Houle, 1992; Falconer and Mackay, 1996; Lynch *et al.*, 1998), however, this expert knowledge is seldom used because it is not clear how to use it in a credible and a consistent manner.

We showed how to use the expert knowledge about the magnitude of different sources of variation by leveraging two recently introduced prior frameworks (Simpson *et al.*, 2017; Fuglstad *et al.*, 2020). While the penalized complexity priors are parsimonious and intuitive, they require absolute prior statements when used in a component-wise approach, which are challenging to elicit for multiple effects. The hierarchical decomposition framework imposes a tree structure according to a domain model, and the intuitive penalized complexity prior can be used in the hierarchical decomposition prior framework to ensure robust modelling. This model-wise approach enables the use of *relative* prior statements, which are less challenging to elicit than the absolute prior statements, because we tend to have good knowledge of the broad sense heritabililty for most traits and by the standard quantitative genetic model construction we know that additive effects capture majority of genetic variance (Hill *et al.*, 2008; Mäki-Tanila and Hill, 2014; Hill and Mäki-Tanila, 2015; Huang and Mackay, 2016). The presented priors therefore pave a way for a fruitful elicitation dialogue between a geneticist and a statistician (Farrow, 2013). In particular, the hierarchical decomposition prior framework provides both a method for prior construction and a platform for communication among geneticists and statisticians. The model-wise expert knowledge prior must naturally be adapted to each model, as it depends on the model structure, but using the tree structures makes this adaption intuitive and should as such help with prior elicitation (O’Hagan *et al.*, 2006; Farrow, 2013). Further, the graphical representation allows a description of a joint prior in a visual way with minimal statistical jargon (Figure 1).

An example of using such expert knowledge was the choices of a median for the broad-sense heritability of 0.25 in the simulated and 0.75 in the real case study. However, as Figures 3 and S6 show, the priors do not differ tremendously. This shows that the prior proposed in this study is vague and not restricted by the value chosen for the median. Perhaps there is even scope for more concentrated priors, should such information be available.

The hierarchical decomposition prior framework enabled us to use expert knowledge on relative additive and non-additive variation. If non-additive effects are to be added to the model, expert knowledge is necessary for the inference to be stable and the results reliable, and the simulation study shows that the expert knowledge must be added in such a way that the magnitude of the variances are not restricted by the prior, i.e., the model-wise approach instead of the component-wise approach. In the simulated case study the expert knowledge improved the stability of inference of the Bayesian approach over the maximum likelihood approach and improved the selection of the genetically best individuals. This improvement was due the additional information that alleviated the strong confounding between the non-additive (particularly epistasis) and environmental variation.

The hierarchical decomposition prior framework is also useful when expert knowledge is only available on parts of the model. For example, an expert may not have a good intuition about the level of broad-sense heritability, say for a new trait, but will likely have a good intuition on how the genetic variance *relatively* decomposes into additive, dominance and epistasis components, simply due to the model specification (Hill *et al.*, 2008; Mäki-Tanila and Hill, 2014; Hill and Mäki-Tanila, 2015; Huang and Mackay, 2016). In those cases, we can use weakly-informative default priors on the parts of the model where expert knowledge is missing, and priors based on expert knowledge for the rest of the model. The component-wise specification of expert knowledge with the standard (Sorensen and Gianola, 2007) or the penalized complexity (Simpson *et al.*, 2017) priors is infeasible in this context, and does not admit a simple visualization of the implied assumptions on the decomposition of the phenotypic variance. Further, the component-wise specification of expert knowledge is particularly challenging when phenotypic variance is unknown or when collected observations are influenced by a range of effects which can inflate sample phenotypic variance. The model-wise approach with the hierarchical decomposition prior can address these situations.

There exists previous work on penalized estimation of genetic covariances (e.g., Meyer *et al.*, 2011; Meyer, 2016, 2019) that also uses Bayesian principles and scale-free penalty functions to reduce variation of the estimates from small datasets and for large numbers of traits. Our proposed priors and expert knowledge reduced variation of estimates in the simulated case study. However, our estimates were biased, which is expected given the small sample size and that the Bayesian approach induced bias towards a lower variance (e.g, Sorensen and Gianola, 2007). It is worth noting that the maximum likelihood estimates of genetic variance also were largely underestimated, which we believe is due to the small sample size and a large number of parameters to estimate. We see in Note S2 in the Supplemental materials (File S1) that the data informs about phenotypic variance and broad-sense heritability, but only weakly about the division of the additive and non-additive, and dominance and epistasis. Further, for some datasets we could not obtain the maximum likelihood estimates, while priors robustified the modelling by penalizing the genetic effects. The real case study also showed that using expert knowledge increases the inference robustness in datasets with a large amount of unobserved phenotypes, and reduces overfitting. We saw this improvement in both the Bayesian approach and the maximum likelihood approach where we used the results from the Bayesian models as initial values for the optimization algorithm. However, the latter approach requires specific expert knowledge on the size of the variances, which in the same way as the component-wise expert knowledge priors, is difficult to elicit from experts in the field. We note, however, that genomic models are inherently misspecified by trying to estimate the effect of causal loci through correlated marker loci (Gianola *et al.*, 2009; *de los Campos et al.*, 2015). Also, linkage and linkage disequilibrium are challenging the decomposition of genetic variance into its components (Gianola *et al.*, 2013; Morota *et al.*, 2014; Morota and Gianola, 2014). Indeed, our variance estimates were not very accurate in the simulated case study.

Future research could expand the hierarchical decomposition prior framework to other settings. For example, to multiple traits or modelling genotype-by-environment interactions, which are notoriously noisy, and we aim to find parsimonious models (e.g., Meyer, 2016, 2019; Tolhurst *et al.*, 2019). Also, expand to model macro- and micro-environmental effects (e.g., Selle *et al.*, 2019) and to model multiple layers of sparse, yet high-dimensional, “omic” data from modern biological experiments using network-like models (Damianou and Lawrence, 2013).

### Importance of priors for breeding

Robust genomic modelling of non-additive variation is important for breeding programs. There is substantial literature indicating sizeable non-additive genetic variation (e.g., Oakey *et al.*, 2006; Muñoz *et al.*, 2014; Bouvet *et al.*, 2016; Varona *et al.*, 2018; de Almeida Filho *et al.*, 2019; Santantonio *et al.*, 2019; Tolhurst *et al.*, 2019), but robust modelling of this variation is often challenging. We have shown how to achieve this robust modelling with the proposed priors and expert knowledge. We evaluated this approach with a simulated wheat breeding program where we assessed the ability to select the genetically best individuals on their genetic value (variety selection) and additive value (parent selection). The results showed that the proposed priors and the expert knowledge improved variety and parent selection. We observed more improvement in the variety selection, which is expected because there is more variation in genetic values than its first-order approximation additive values. However, this additional non-additive variation is hard to model due to a small signal from the data relative to environmental variation and confounding with the environmental variation. This confounding is expected. As pointed by one of the reviewers, we obtain the epistasis covariance matrix using the Hadamard product of the additive covariance matrix with itself, and such repeated Hadamard multiplication converges to an identity matrix, i.e., to the covariance matrix of the environmental effect. Both the simulated and real case studies showed that including non-additive effects in the model requires some sort of penalization to avoid overfitting environmental noise as non-additive genetic effects. The proposed priors and the expert knowledge helped us to achieve this.

Importantly, all models improved upon sole phenotypic selection in the simulated case study, which shows the overall importance of genomic modelling While the differences between the different models and priors were small, the improved genomic modelling can contribute to the much needed improvements in plant breeding (Ray *et al.*, 2013; *Asseng et al.*, 2015). Also, even a small improvement in the variety selection has a huge impact on production, because varieties are used extensively (Acquaah, 2009). In the terms of model complexity, the answer to whether to use the additive model, the additive and dominance model or the non-additive model depended on the aim of the analysis. The latter models were the best in selecting the genetically best individuals on genetic value, whereas the additive model performed best in selecting the genetically best individuals on additive value. The reason for this is likely the small sample size and large number of parameters to estimate with the non-additive model (Varona *et al.*, 2018). In the real case study adding non-additive effects to the model improved the phenotypic prediction accuracy beyond the additive model, and the expert knowledge helped us to avoid overfitting, which shows the advantage of the expert knowledge.

Of note is the observation that the proposed priors and the expert knowledge improved the selection of the genetically best individuals, but not the estimation of the different genetic values. We did not expect this difference. In principle both of these metrics are important, but for breeding the ability to select the genetically best individuals is more important (de los Campos *et al.*, 2013). A possible explanation for the difference between the two metrics is that the top individuals deviated more from the overall distribution and the overall metrics do not capture well the tail-behaviour.

The importance of the proposed priors and the expert knowledge will likely vary with the stage and size of a breeding program, and as the simulation study with increasing amount of observations and the real case study shows, the importance of priors increases with the decreasing amount of observations. Prior importance is known to decrease as the amount of data increases (Sorensen and Gianola, 2007), but the required amount of data for accurate estimation of non-additive effects is huge compared to the size of most breeding programs. Therefore the proposed penalized complexity and hierarchical decomposition priors could be helpful also in large breeding programs as they enforce shrinkage according to the expert knowledge unless the data indicates otherwise, reducing the risk of estimating spurious effects.

### Limitations of our work

The aim of this paper was to describe the use of the expert knowledge to improve genomic modelling, which we achieved through two recently introduced prior frameworks (Simpson *et al.*, 2017; Fuglstad *et al.*, 2020), and demonstrated their use in a simulated and a real case study of wheat breeding. There are multiple other possible uses of the proposed priors in genomic modelling and prediction. The simulated case study is small with only 100 individuals at the advanced yield trials of a wheat breeding program, and up to 700 individuals at the preliminary yield trials. A small number of individuals and a limited genetic variation at this stage made a good case study to test the importance of priors, and we show that using our approach can be beneficial beyond the standard genomic model. We have also chosen this stage for computationally simplicity and speed because we evaluate the robustness of estimation over many replicates. Studies with more individuals are a natural next step, but is beyond the scope of this paper due to computational reasons. Finally, we could have tested more prior options, in particular the shrinkage of the non-additive values towards the additive values, i.e., the PC_0_(·) versus the PC_M_(·) prior. More research is needed in the future to see how the expert knowledge can improve genetic modelling further.

Interesting areas for future research are also in other breeding domains with the recent rise in volumes of individual genotype and phenotype data, which provide power for estimating dominance and epistasis values (e.g., Alves *et al.*, 2020; Joshi *et al.*, 2020). The ability to estimate the non-additive values would be very beneficial in breeding programs that aim to exploit biotechnology (e.g., Gottardo *et al.*, 2019). Finally, an exciting area for estimating non-additive individual values is in the area of personalized human medicine (de los Campos *et al.*, 2010; Mackay and Moore, 2014; Sackton and Hartl, 2016; de los Campos *et al.*, 2018; Begum, 2019).

The proposed priors are novel and require further computational work to facilitate widespread use. The penalized complexity priors (Simpson *et al.*, 2017) are increasingly used in the R-INLA software (Rue *et al.*, 2009, 2017), while hierarchical decomposition priors (Fuglstad *et al.*, 2020) have been implemented with the general purpose Bayesian software Stan (Carpenter *et al.*, 2017; Stan Development Team, 2019). This implementation is technical and Stan is slow for genomic models, although there is active development to increase its computational performance (Margossian *et al.*, 2020).

We are in the process of developing an R package that will offer an intuitive user-interface to specify hierarchical decomposition priors. The clear graphical representation of the priors along the model defined tree will encourage increased transparency within the scientific community. It will facilitate communication and discussion between statisticians and non-statisticians in the process of the model design, prior specification but also model assessment. Existing expert knowledge is intuitively incorporated into PC prior distributions for the parameters where it applies to. The resulting model-wise prior can be fed directly into Stan or INLA, or can be pre-computed for use in other Bayesian software. Thus, the new priors will be straightforward to apply for statisticians and non-statisticians, robustify the analysis, and the use of INLA will speed up computations. Further work is needed to enable Bayesian treatment of large genomic models fitted to datasets with hundreds of thousands of individuals.

## Conclusion

In conclusion, we have presented how to use the expert knowledge on relative magnitude of genetic variation and its additive and non-additive components in the context of a Bayesian approach with two novel prior frameworks. We believe that when modelling a phenomenon for which there exists a lot of knowledge, we should employ methods that are able to take advantage of this resource. We have demonstrated with a simulated and a real case study that such methods are important and helpful in the breeding context, and they might have potential also in other areas that use genomic modelling.

## Acknowledgments

Hem, Fuglstad and Riebler were supported by project number 240873 from the Research Council of Norway. Gregor Gorjanc acknowledges support from the BBSRC to The Roslin Institute (BBS/E/D/30002275) and The University of Edinburgh’s Data-Driven Innovation Chancellor’s fellowship.

## Footnotes

a Additive model with component-wise expert knowledge prior.

b Additive model with component-wise default prior.

a Additive and non-additive model with model-wise expert knowledge prior.

a Additive and dominance model with model-wise expert knowledge prior.

a Additive model with model-wise expert knowledge prior.

b Additive model with model-wise default prior.

